# Extensive global HIV-1 sequence diversity perturbs conserved features of the envelope glycan shield

**DOI:** 10.64898/2026.08.20.745985

**Authors:** Maddy L. Newby, Joel D. Allen, Emily R. Lucas, Arturo Monzon, Nicole E. James, Katie J. Doores, Max Crispin

## Abstract

The extensive sequence diversity and glycosylation of the HIV-1 envelope glycoprotein pose a challenge for vaccine design efforts aimed at eliciting glycan-binding broadly neutralizing antibodies (bnAbs). Understanding how HIV-1 glycosylation varies across diverse strains is therefore of considerable interest. Here, we employed mass spectrometry to map the site-specific glycan composition of pseudoviruses from an established 12-virus panel representing global sequence diversity. We refine the current model of the viral glycan shield by showing how the presence or absence of glycans can modulate the composition and perimeter of the ‘mannose patch’, where glycans exhibit limited maturation. Importantly, we show that within the Clade A 398F1 strain, isolated in Tanzania, the trimer apex exhibited elevated glycan maturation and resistance to apex-directed bnAbs. These findings demonstrate that the 12-virus panel reveals both conserved glycan-dependent epitopes and strains with exceptional glycosylation features that may impact the generality of vaccine design efforts targeting particular epitopes.

**Highlights:**

- Global HIV-1 diversity preserves key features of the envelope glycan shield
- Glycan occupancy and local networks reshape the perimeter of the mannose patch
- 398F1 displays atypical apex glycosylation linked to bnAb resistance
- Some glycan-dependent bnAb epitopes remain robust across diverse HIV-1 strains

**In Brief:** Newby et al. map site-specific glycosylation across a globally representative 12-virus HIV-1 panel. Despite extensive sequence diversity, several glycan-dependent bnAb epitopes are conserved, while strain-specific differences in glycan occupancy and processing reveal features that may contribute to neutralization resistance.

## Introduction

Recent clinical trials have demonstrated a pathway towards the generation of a protective HIV-1 vaccine through an approach aimed at eliciting broadly neutralizing antibodies (bnAbs)^1–4^. Antibody breadth is a critical parameter for HIV-1 vaccine development due to the immense diversity of circulating viruses^5,6^. BnAbs with exceptional breadth and potency are protective in challenge studies in non-human primates and have been shown to protect against infection of sensitive viruses in humans^7–9^. Multi-stage strategies to guide antibody affinity maturation specifically towards bnAb elicitation remain a central goal of HIV-1 vaccine design^10–13^.

The HIV-1 Envelope glycoprotein (Env), comprising a trimer of gp120-gp41 heterodimeric protomers, is the sole viral target of bnAbs and is extensively shielded by host-derived N-linked glycans. A single Env protomer contains around 24-30 N-linked glycan attachment sites (encoded by NxS/T, where x ≠ P), representing one of the most densely N-glycosylated proteins in nature. As these glycans are host-derived, antibody responses against them are often limited by immune tolerance mechanisms that suppress autoreactive B cells^14^, especially when compared to the antigenic protein surface that is covered by the glycan shield^15^. Despite these challenges, almost all broad and potent bnAbs recognize or accommodate specific HIV-1 Env glycans as part of their epitopes^16–18^, afforded by one or multiple of the defined features of bnAbs such as long heavy-chain complementarity determining regions (CDRs), rare germline usage, and high somatic hypermutation (SHM)^19–21^. In-depth understanding of the molecular components of a bnAb epitope is valuable for rational vaccine design^22^.

Key bnAb epitopes on Env include the CD4-binding site (CD4bs), V1/V2 apex, V3 loop, gp120–gp41 interface, the gp120 silent face (SF), and the membrane proximal external region (MPER)^23,24^, with the first five regions comprising heavily of glycans^18,25^. Potent bnAbs target Env through a combination of protein-protein and protein-glycan contacts, with the latter possible due to the conserved nature of key underprocessed glycans on Env^26,27^. Although N-linked glycan attachment occurs co-translationally, subsequent processing continues as the glycoprotein egresses through the endoplasmic reticulum and Golgi apparatus. The processing stages that occur immediately following protein folding are influenced by the tertiary and quaternary architecture of Env^28,29^. The dense steric environment of the Env glycan shield limits the ability of glycan processing enzymes to fully mature Env N-linked glycans, resulting in a predominance of underprocessed oligomannose-type glycans rather than the heterogeneous mixture of complex-type glycans typical of hostglycoproteins^26,27,30,31^. These types of glycans are recognized by a range of bnAb classes, most notably V3 glycan, V2-apex, and SF bnAbs^32,33^.

Despite their extraordinary breadth and potency, bnAb resistance remains a significant therapeutic threat, as underscored by the results from the Antibody Mediated Prevention (AMP) trial where only one third of existing circulating viruses were adequately sensitive to the CD4bs-directed bnAb VRC01 for prevention at the doses tested in the AMP trials^34,35^. In-depth information exists on bnAb resistance, ranging from specific point mutations and increasing variable loop length, to the modulation of N-linked glycan sites^36–38^. Although protein structure has emerged as an important parameter in shaping glycan maturation, glycosylation is often non-templated beyond the initial N-linked glycosylation attachment sequon, and the sequon alone does not directly dictate the resultant glycan composition that is present on mature viruses. Therefore, the composition of Env glycosylation cannot be readily deduced from nucleotide sequence alone, requiring orthogonal analytical approaches. These approaches are important in vaccine design given the key roles of glycans in viral diversification and immune evasion.

Since wide antibody breadth and high potency are key attributes for establishing effective correlates of protection against HIV-1, extensive research has led to the development of virus panels that capture the global diversity of HIV-1, including the widely used 12-virus HIV-1 panel described by deCamp, A., *et al.*^39^. The Env strains in this panel represent a broad neutralization profile encompassing multiple major genetic clades and circulating recombinant forms, including clade A (398F1), B (X2278 and TRO11), C (CE0217, CE1176, and 25710), G (X1632), CRF01_AE (CNE8 and CNE55), CRF07_BC(CH119 and BJOX002000), and AC recombinant (246F3). The widespread use of this panel has generated a wealth of bnAb sensitivity data, positioning it as a valuable resource. However, despite the role of glycosylation in bnAb neutralization and resistance, only limited information is available on the composition of the glycan shield from HIV-1 virions^26,27,40,41^, and knowledge across this virus panel remains particularly poorly understood. Soluble recombinant formats of Env, including trimeric gp140 and monomeric gp120, have historically provided a route to understand HIV-1 Env glycosylation and have led to the characterization of key glycan-based bnAb epitopes. However, these recombinant formats are limited and tend to accentuate non-native features of Env glycosylation, particularly at the trimer apex and surrounding the CD4bs^29,42,43^. To unpick the potential glycan-mediated mechanisms of sensitivity and resistance to neutralization by bnAbs, it is therefore necessary to capture the glycosylation of native-like Env trimers across genetically diverse strains using a scalable and broadly applicable expression platform. To address this, we analyzed the glycosylation of Env isolated from pseudoviruses comprising the global 12-virus panel and propose potential glycan-mediated mechanisms of bnAb resistance. These data provide additional molecular context for the interpretation of neutralization profiles of these strains against diverse nAbs.

Using liquid chromatography-mass spectrometry (LC–MS), we analyzed the site-specific glycosylation of all 12 pseudovirion-derived Envs in this panel, performing several independent expressions for each strain. This analysis revealed remarkable conservation in glycan processing at sites key for bnAb binding such as N160 and N332, in addition to strain-specific variation in glycan processing that are attributed to the surrounding local glycan networks and protein architecture. Furthermore, we identify potential N-linked glycan sites (PNGS) that are either highly underoccupied or completely unoccupied by glycans, with a particular enrichment in the V1/V2 loops. These data refine our understanding of the viral glycan shield and will provide an important resource to the HIV-1 vaccine community, aiding an understanding of how glycobiology contributes to bnAb resistance. Crucially, for vaccine design efforts we demonstrate both the robustness of particular regions of the glycan shield that form a focus for bnAb-eliciting vaccines, together with exceptions where glycosylation features are notably poorly conserved.

## Results and Discussion

### N-linked glycosylation sequons are abundant across Env, with key hotspots

As extensive glycosylation is observed on the Env of every HIV-1 strain, we sought to understand the variation in location and frequency of N-linked glycan attachment sites. To do this, we utilized all available sequences from the CATNAP database ranging from 1983– 2024, comprising over 2400 sequences, and determined the HxB2 aligned glycosylation positions using the ‘N-glycosite’ tool^44,45^. Across all sequences, the number of N-linked glycosylation sites in a single Env protomer ranged from 18 sites on clade A isolate 93UG077 AY669704 to 37 on clade B isolate N90 08B6.JQ610142, with an average of 29 sites across all samples (**Figure 1A** and **Data S1**). Across clades, the overall number of sites was broadly comparable, with the mean number of sites within clades A, B, C, D, F, G, CRF01_AE, CRF02_AG, and CRF07_BC ranging from 28-30 sites per protomer (**Figure 1B**). When comparing the location of the N-linked glycosylation attachment sequons, two main trends emerged. Firstly, of the 283 unique HxB2-aligned N-linked glycosylation positions, only 17 were conserved in 75% of sequences and, of these, 9 were conserved in 95% of sequences: N88, N156, N197, N241, N262, N301, N611, N625, N637 (**Figure 1C** and **Data S1**). In contrast, 195 unique N-linked glycosylation positions were observed in <5% of sequences. This highlights the duality of the Env glycan shield; whilst there are regions that are highly conserved and likely critical for immune escape or glycoprotein function, most sites are highly variable. These variable sites tend to be located within the variable loops (**Figure 1D-E**), with V1/V2 sampling 83 different locations alone.

**Figure 1.**
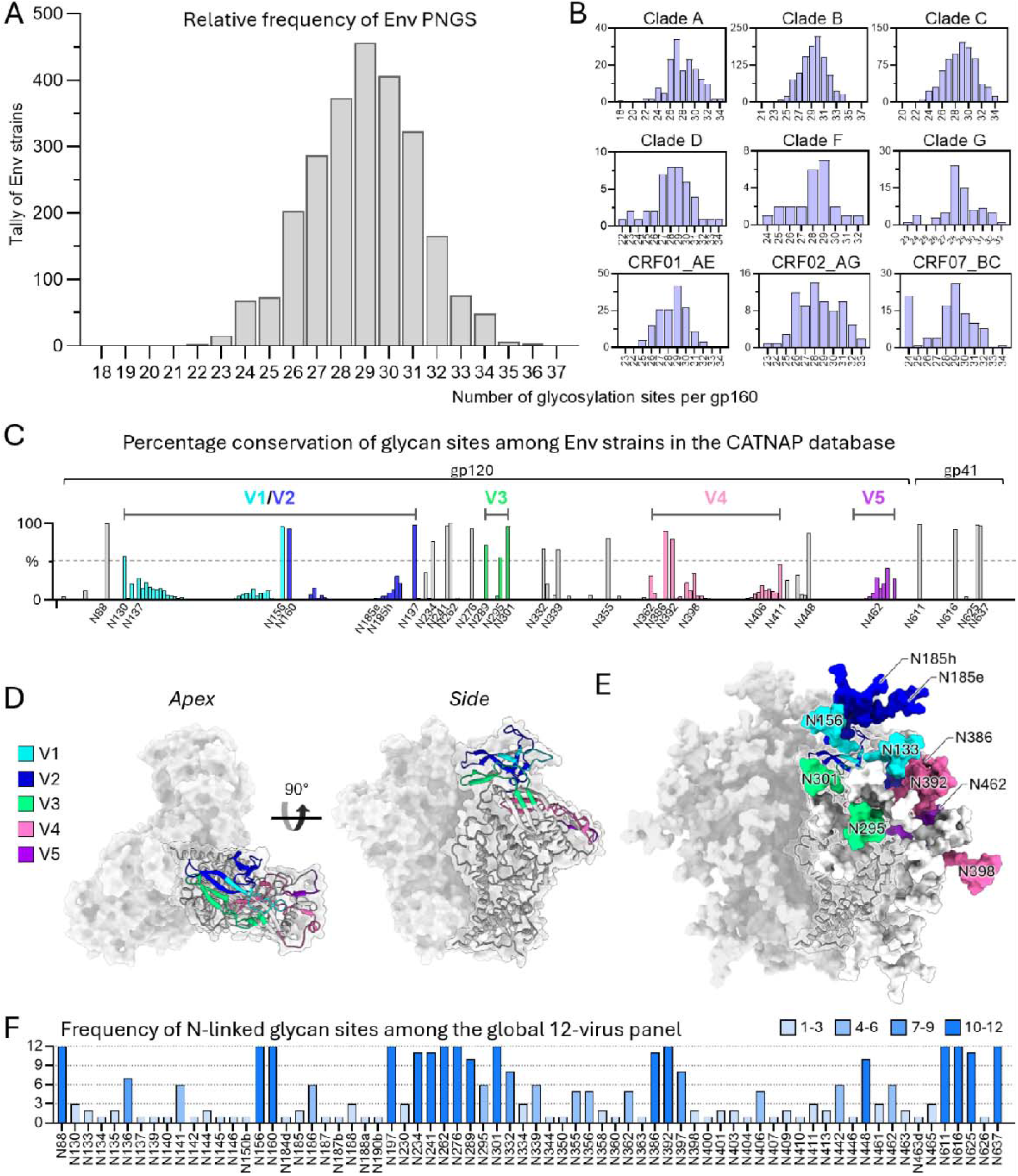
Histograms indicating the relative frequency of N-linked glycan sites on Env. **A)** Histogram displaying the relative frequency of PNGS on gp160 (PNGS up to position N648) from all group M HIV-1 Env strains deposited on the HIV database (www.hiv.lanl.gov, 2024 listing^44^). **B)** Data from (A) is further extrapolated according to the clades from which the Env strain originates to display the clade-specific distribution of PNGS. **C)** The frequency of PNGS across gp160 with the PNGS found in the reference BG505.T332N strain labeled on the X axis. **D)** Structure of the Env trimer using PDB: 5ACO showing the apex (left) and side (right) orientations. Two protomers are shown in the surface representation and one shown in ribbon with variable loops distinctly colored according to panel C. **E)** Glycosylated Env model with variable loop glycans colored according to panel C and labeled with the HxB2-aligned Asn residue. **F)** Conservation of N-linked glycan sites across the global 12-virus panel. The frequency of PNGS is colored according to the extent of conservation within the virus panel, ranging from light blue (representing low conservation) to dark blue (representing high conservation).

Outside of the nine highly conserved glycosylation sites, clade-specific variation was observed. One example is the N130 glycan, which has been repeatedly shown to limit V2 apex bnAb neutralization breadth^46–49^. In the CATNAP database, among clades with >20 available sequences, the highest occurrence of N130 is in clade CRF01_AE (70% of 159 strains), while in clade A it is present in 31% of sequences. Despite expected variations in the frequency of glycosylation sites within the variable region, there are clade-specific differences in the presence of more conserved glycosylation sites. Glycans at N230, N234 and N241 can influence CD4bs bnAb neutralization and display clade-specific variation. The N234 site is present in only 66% of clade B sequences but occurs in 85% of clade A sequences. The N230 glycan is much less conserved overall, yet appears in 75% of clade C sequences compared to 29% in clade B. In contrast, N241 is highly conserved, present in 85% of clade A sequences, and its absence has been shown in some contexts to elicit autologous neutralizing antibodies^50,51^.

The N295 glycan, which contributes to both V3 glycan and SF bnAb epitopes, also demonstrates clade-specific variability. This glycosylation site is present in only 22% of clade C sequences and 2% of CRF07_BC sequences, compared to 73% of clade B sequences. Likewise, compared with other regions of the glycan shield, the abundance of N332 versus N334 glycosylation sites vary. The mechanism by which N332 shifts to N334 to escape V3 glycan neutralization is well described^6,36,38,52,53^. The N332 site was observed within 75% of sequences within clade C and N334 is observed 14% of the time. Whereas, within CRF01_AE clade, the most abundant circulating clade in southeast Asia, this observation is reversed, with N332 in 12% of sequences and N334 in 84% of sequences. This demonstrates the pervasiveness of this glycan escape mutation across the global Env populations.

Other interesting observations include the presence of N363 in >50% of clade A and CRF02_AG but <15% in other well-represented clades, and the absence of the N160 glycan, which is often critical for V2 apex neutralization, in 16% of clade G sequences. Lastly the N276 glycan site is moderately underrepresented within CRF07_BC (∼70% of sequences) compared to 85% across other clades. This glycan hinders antibody access to the CD4bs, and its removal promotes germline CD4bs antibody binding^12,54–56^.

The global 12-virus panel was previously curated by deCamp *et al.* with the consideration of capturing the major patterns of antibody-mediated neutralization observed across circulating HIV-1 strains^39^. Overall, these viruses sample 72 unique N-glycosylation sites (**Figure 1F**) and include viruses that represent the dominant V3 glycan cluster configurations, with either N332 (present in ∼70–75% of global isolates) or its escape variant N334 (∼15–20%). Similarly, V2 apex glycans N156 and N160, together with N197, N276, and N301, are highly conserved (>70% prevalence globally) and are well represented in the panel. The N295 glycan is also represented at a frequency representative of global variation. Overall, the global 12-virus panel mirrors the conservation and variability of glycan sites observed in global Env populations, incorporating both highly conserved glycosylation and positional variation. Whilst the location and conservation of glycan sequons play an important role in understanding bnAb neutralization, the processing state of the attached glycan plays a central role. Further, the presence or absence of particular glycan sites influences the attachment and processing of surrounding sites. Additional analyses are required to understand how perturbations in the glycan shield can influence antibody binding^31,43^.

### Mannose glycans persist in the face of sequence diversity

To determine the composition of the glycan shield of the global 12-virus panel, we utilized a third-generation lentivirus packaging system to produce non-replicating HIV-1 pseudoviruses^57^. The secreted pseudoviruses were purified using *Galanthus nivalis* lectin (GNL), which recognizes the α-1,3-linked mannose that is abundant on the oligomannose-rich HIV glycan shield and ostensibly captures virions from across the full viral pool. Independent preparations of each pseudovirus-derived Env were proteolytically digested and analyzed by LC–MS to characterize the glycan shield at the site-specific level.

Using this approach, we were able to obtain glycosylation data for 326 individual sites, covering 71 of the 72 unique glycosylation sites across the 12 viruses (**Figure 2A**). Across all Envs analyzed in this study, the most striking feature is the pervasiveness across gp120 of high-mannose glycans (which is used here for methodological reasons to refer to endoglycosidase H (Endo H)-sensitive glycans that include both oligomannose- and hybrid-type glycans). Out of the 71 sites analyzed, 32 contained >50% high-mannose glycans and 19 sites contain ≥90% high-mannose glycans (**Figure 2A-B**). The extensive display of high-mannose glycans has been reported previously on a handful of strains but is also heavily reflected across the diverse global 12-virus panel^26,27^. In particular, glycan sites located at protomer interfaces and on the outer domain of gp120 are hotspots for high-mannose glycans, and these sites are not only limited to the conserved regions of Env but are also observed in the variable loops. Of the 9 highly conserved sites within the CATNAP database (>95% conservation), N156, N262, and N241 were consistently dominated by high-mannose glycans across all strains in the global 12-virus panel. Of note, N156 and N262 were occupied by an average of 97.4 and 98.4% high-mannose glycans, respectively (normalized against PNGS occupancy). These sites are representative of the two distinct mannose patches located on Env; the so-called intrinsic mannose patch (IMP; includes N262) and the trimer associated mannose patch (TAMP; includes N156^29^). The TAMP represents PNGS that are primarily occupied by high-mannose glycans only when Env is assembled as a fully cleaved trimer^28^. Therefore, these data demonstrate that Envs isolated from pseudovirions adopt a native-like quaternary architecture. Outside of these regions, the high-mannose glycan content is lower and much more variable, particularly within the gp120 variable loops and on gp41 (**Figure 2C**).

**Figure 2.**
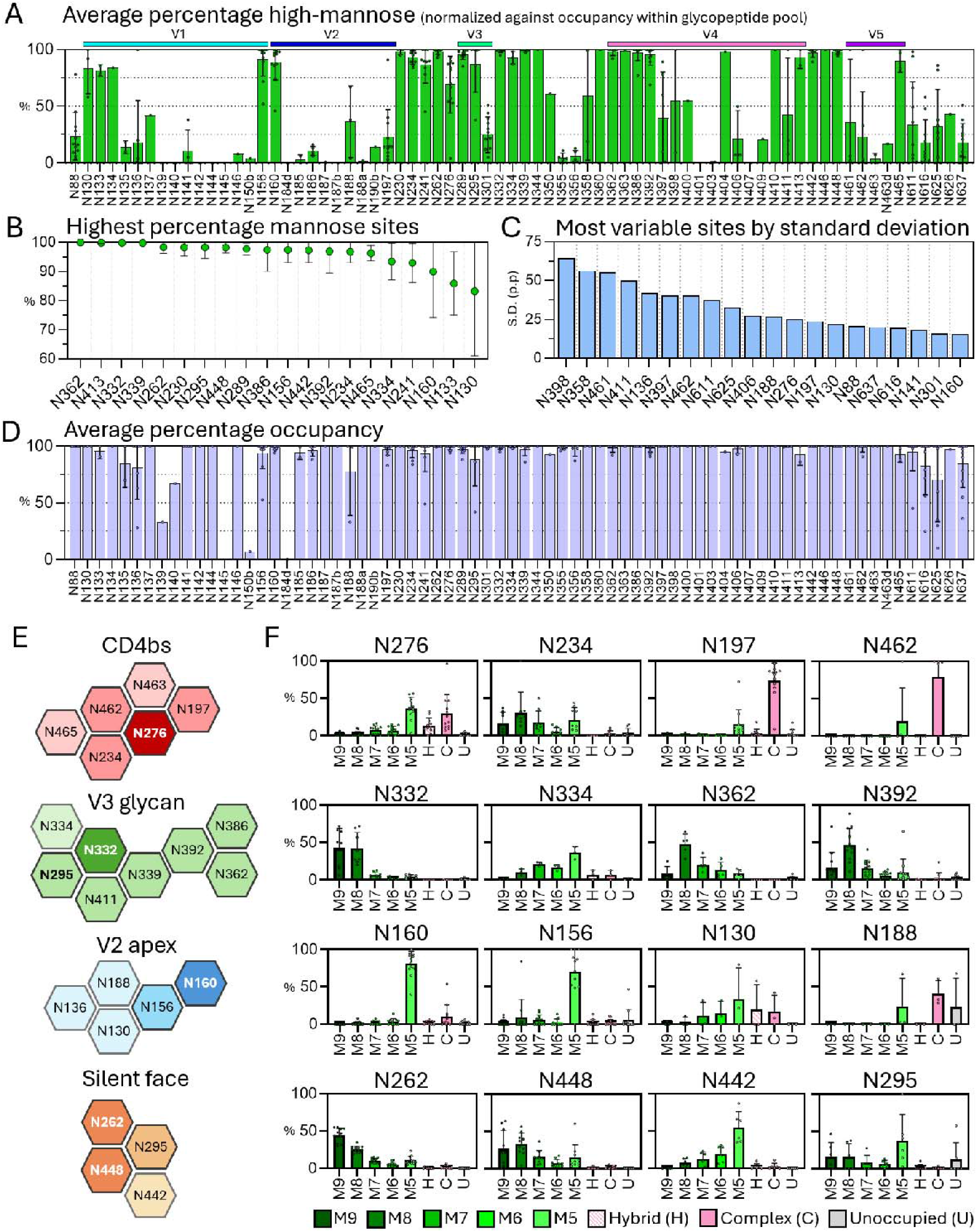
Average epitope-specific glycosylation. **A)** The abundance of high-mannose glycosylation at unique PNGS within the 12-virus panel, normalized against occupancy. Individual points represent the percentage abundance for each Env strain (averaged across multiple independent expressions), with each bar representing the average abundance across all Env strains containing that site. Variable loops are annotated above the graph using colors used in Figure 1. **B)** Glycan sites ranked in order of highest abundance of oligomannose-type glycosylation. Only sites that are present in ≥2 strains are included. **C)** PNGS exhibiting the largest variation in glycan processing as a measure of standard deviation (S.D.), with glycosylation data normalized against PNGS occupancy. **D)** Average site-specific PNGS occupancy. Individual data points represent PNGS occupancy averaged across independent expressions for each Env strain. **E)** The network of glycans surrounding the CD4bs, V3-glycan, V2 apex, and silent face epitopes. The glycans that comprise the central bnAb target for each epitope are depicted using a brighter color. Glycans that are more peripheral and form fewer contacts with bnAbs are colored in decreasing saturation. **F)** Average fine processing of glycans implicated in bnAb recognition across every strain in the global 12-virus panel. The average relative percentage abundance of each glycan type across all 12 strains is plotted as a single point, and the average glycosylation across the panel of pseudoviruses are represented as bars. The glycan categories are split into Man_9_GlcNAc_2_ – Man_5_GlcNAc_2_ (M9–M5); Hybrid (H); complex (C); and unoccupied (U). In all graphs, the bar represents the average value across the global 12-virus panel and error bars represent the standard deviation (S.D.).

In addition to identifying sites where glycan processing is highly conserved across the virus panel, this analysis also highlights site-specific deviations. Several sites exhibit substantial variation in glycan processing across strains, reflected by large standard deviations presented in **Figure 2A** and **C**, and are visualized in **Figure 3**. Two such examples include N358 and N461, where a shift in glycan processing from complex- to oligomannose-type can be attributed to the presence of a neighboring glycan that creates a more sterically restrictive environment, causing glycan processing to stall. In the CE1176 strain, the presence of N362 is associated with limited glycan processing at N358, whereas the absence of N362 in X2278 is consistent with full processing at N358. Localized glycan clusters are also observed within the V5 region on 398F1 where glycans at N465 limit extensive glycan processing at N461. Given that conserved contacts with V5 loop residues 456-459 are commonly made by CD4bs bnAbs and the deletion of V5 loop glycans can enhance bnAb neutralization potency^12^, modulation of V5 loop glycosylation could act as a mechanism to impart bnAb neutralization resistance. So, while these analyses identify highly conserved signatures of glycosylation that persist in the face of extensive sequence diversity, variation in glycan processing at the site-specific level can occur due to natural differences in strain- or clade-specific glycan networks.

**Figure 3.**
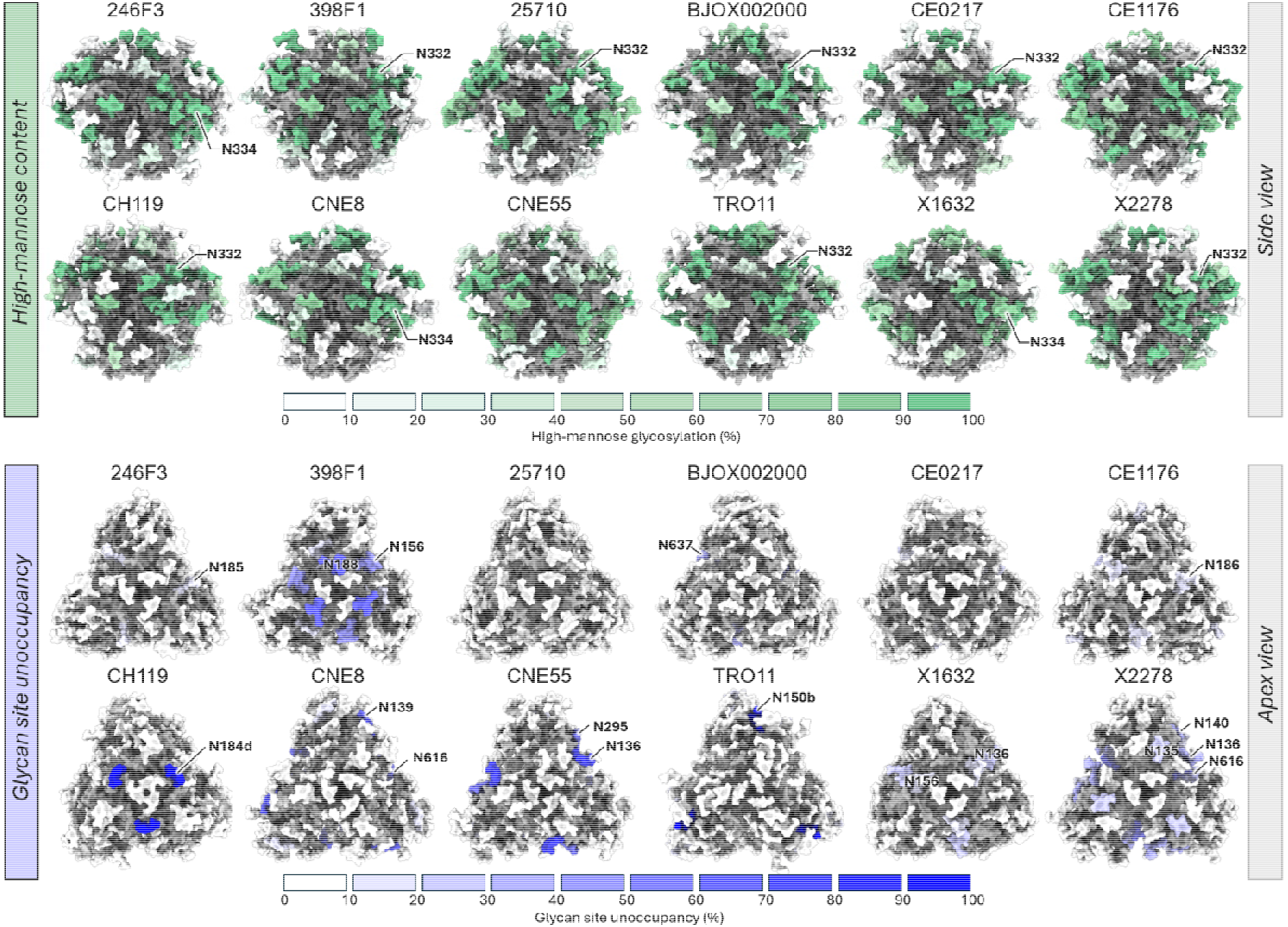
Glycan underprocessing and underoccupancy across the global 12-virus panel. Side and apex views of the AlphaFold3^69^ and GlycoShape Re-Glyco^70^ derived models of the global 12-virus panel Envs. Site-specific N-linked glycan high-mannose quantifications are colored according to the green key, with the N332 supersite or N334 glycans labeled, while site-specific glycan site underoccupancy are colored according to the blue key. N-linked glycan sites displaying >10% unoccupancy are labeled.

### Glycan site underoccupancy is enriched in the V1/V2 loops

With exceptions, high PNGS occupancy was observed across the gp120 region of pseudovirus-derived Env (**Figure 2D**). Previous studies have demonstrated that incomplete glycan occupancy on soluble recombinant Env immunogens converges to near-complete in a virus-like context, with a particular focus on BG505 and a select few other well-characterized Env strains^26,27^. Our analysis is consistent with these previous observations, but here we extend this across a diverse set of Env strains. Previous site-specific glycan analyses have suggested that virus-like Env can have a largely complete glycan shield, whereas our data show that the presence of a PNGS does not always result in glycan occupancy^26,27,58^.

Where Env strains encode two closely spaced PNGS (separated by ≤1 amino acid), simultaneous occupancy may be constrained. We frequently observed mutually exclusive glycan occupancy at these sites (**Figure 2D**). To this end, we observed PNGS that were highly underoccupied (∼5-30% occupancy) or completely unoccupied on TRO11, CNE8, and CH119. Notably, these sites were particularly enriched in the V1/V2 region and include N150b in TRO11, N139 in CNE8, and N184d on CH119 (HxB2 aligned, **Figure 3**). This leads to questions surrounding the role of such PNGS during Env evolution. Whether these preferentially unoccupied sequons arise stochastically or are positively selected due to the biochemical properties of the surrounding primary amino acid sequence is unclear. While the observation of increasing V1/V2 PNGS frequency is well-documented in both cross-sectional and longitudinal studies^36,59,60^, our analyses caution against assuming complete glycan occupancy at all acquired PNGS. A similar observation was observed on the SARS-CoV-2 Gamma (P.1) spike glycoprotein, where a PNGS insertion at N20 ablated glycosylation at the conserved N17 site^61,62^. In perhaps a more extensive example, the V1 sites N135, N136, and N140 (HxB2 aligned; or N134, N137, and N141 in the primary sequence) on X2278 are occupied between 30-38% of the time, suggesting that glycosylation at these sites is mutually exclusive in a different capacity to that described above. A similar distribution in PNGS occupancy has been reported on a variety of human glycoproteins^63–66^.

Outside of the V1/V2 loop, we detected the greatest variation in glycan site occupancy in the gp41 subunit. Evidence for partial glycan occupancy on pseudoviral gp41 has previously been inferred through protein laddering on SDS-PAGE^67^. Here, we have been able to localize this underoccupancy at the site-specific level across the global 12-virus panel. Interestingly, relative to the other gp41 sites, N625 seems to be the most frequently under-occupied, likely due to an unfavorable amino acid sequence context for glycan attachment^68^. Unlike soluble Env trimers, which commonly display an artificial N611 glycan hole^51^, glycan occupancy at N611 averaged at 94.8% across the panel of HIV-1 pseudoviruses. The variation in glycan site occupancy across the panel highlights the fact that incomplete glycan attachment on pseudovirus-derived Env likely occurs in a sequence context-specific manner.

### Glycan heterogeneity defines the CD4bs epitope across global strains

As mentioned, Env glycans often form integral components of bnAb epitopes or can be accommodated to make critical peptide contacts. The glycan networks that comprise, and are involved in the recognition of, bnAb epitopes are schematically depicted in **Figure 2E**.

The CD4bs bnAbs (especially those of the VRC01 and IOMA class) are not dependent on glycans for epitope binding and virus neutralization but instead have evolved to accommodate glycans at N276. These analyses highlight the challenges posed by the N276 glycan in the pathway to inducing CD4bs bnAbs. Whereas sites such as N332 are consistently occupied by oligomannose-type glycans, N276 always contains a mixture of oligomannose-, hybrid-, and complex-type glycans, representing extensive glycan microheterogeneity across the Env population (**Figure 2F**). Furthermore, the ratio of high-mannose to complex-type glycans varies between samples. The lowest high-mannose glycan content at N276 is 3.7% on X1632 and the highest is 92.6% on 246F3, averaging 68.7% across all strains. Another interesting observation is low fucosylation of complex-type glycans at N276, with only an average of 17.2% of glycans containing fucose at this site (**Data S2**).

Surrounding the CD4bs epitope lie glycans at N197, N234, and those in the V5 loop (**Figure 2E**, top). While not directly implicated in bnAb binding, these glycans represent a barrier to CD4bs bnAb induction, substantiated by the fact that germline targeting strategies remove these glycans to promote germline VRC01-class antibody binding^71^. The N234 glycan site is consistently occupied by high mannose-type glycans across all analyzed Env strains. The opposite is true for glycans at N197, which, across all strains except BJOX002000, are efficiently processed to complex-type. Out of the panel viruses, BJOX002000 is the most resistant to the CD4bs bnAbs VRC01, IOMA, and 3BNC117 (geometric mean IC_50_). Extensive mannosylation at both N234 and N197 on BJOX002000 (89.4% and 90.9%, respectively) compared to that of the other global panel viruses (average: 93.4% and 17.0%, respectively) could indicate that the CD4bs epitope is more sterically restricted in this strain (**Data S2**). The V5 loop PNGS across the panel viruses are diverse in location (**Figure 1C**). Despite this, complex-type glycans persist across all but three (398F1, CNE55, and CNE8) panel pseudoviruses. From these analyses, patterns in V5 loop glycosylation are apparent: the presence of a single V5 glycan site allows for full glycan maturation presumably due to the enzyme-accessible nature of these glycans, whereas the presence of two glycan sites leads to restricted processing at both sites (**Figure 2A**).

### Microheterogeneity of oligomannose-type glycans within the V3 eptiope

In contrast to the CD4bs, the V3 epitope consists of glycan sites that are critical for high potency virus neutralization by most V3 bnAbs. The N332 site, which is preserved amongst ∼70-75% of circulating viruses (**Figure 1C**), is abundantly occupied by Man_9-8_GlcNAc_2_ (M9-M8) glycans in all analyzed strains containing this site (n = 8) (**Figure 2F**). This demonstrates the resilient nature of this glycan epitope across strains from a range of clades when in the context of the surrounding glycan network. This can be further illustrated by the fact that a positional shift from N332 to N334 has the capacity to induce a subtle yet consistent increase to the extent of mannose trimming at this site^59^. The N362 site forms a part of the IMP microcluster on the outer domain of gp120. Of the strains within the panel that possess this site (n = 5), the degree of mannose trimming is remarkably conserved across the different Env strains, with M8 glycans most abundant in all instances (**Figure 2F**). These data further exemplifies how selective evolutionary constraints preserve key glycan signatures for glycan-dependent bnAb recognition. Conversely, glycan processing at N392 varies slightly more relative to N332 and N362. While still predominantly a ‘mannose’ site, greater microheterogeneity can be observed across the strains that contain this site, likely reflecting its position on the boundary of the IMP. So, while it is still structurally and functionally linked to the V3 glycan epitope, the processing state of this site is likely more extensively influenced by the local architecture.

### Glycan heterogeneity surrounds key V2-apex oligomannose-type glycans

The N160 glycan is central to the binding modality of many V2 apex bnAbs including PG9, PG16, PGT145, and PGDM1400^72–74^. Across the panel viruses, glycan occupancy of N160 is near-complete and high mannose-type glycosylation highly prevalent, with an average of 98.7% and 89.1%, respectively (**Figure 2A** and **D**, **Data S2**). Binding studies using BG505 SOSIP.664 alongside virus neutralization assays have demonstrated that smaller high mannose-type glycans favor high affinity binding and high potency neutralization by V2 apex bnAbs^73,74^. The smaller size of a Man_5_GlcNAc_2_ (M5) glycan compared to larger M9-8 glycans are favorable for CDRH3 accessibility to the trimer apex. The identification of M5 at N160 across the majority of the panel viruses (except for 398F1, which is expanded upon in the following section) provides reassurance that the carbohydrate element of this epitope is well-conserved and targetable across different geographical clades. While not a primary glycan contact, V2 apex bnAbs can make limited contacts with glycans at N156. This site is primarily modified by oligomannose-type glycans, but even hybrid-type glycans at N156 can be tolerated by bnAbs^75^. Like those occupying N160, the predominant glycoform we identified at N156 is M5, however glycans at N156 display more extensive microheterogeneity compared to N160 (**Figure 2F**).

Certain glycans situated at the perimeter of the V2 apex epitope have been shown to hinder bnAb binding and virus neutralization^46–49,73^. Consistent with the highly variable nature of the V1/V2 loops in terms of both the position and number of PNGS, we observed variable glycosylation in this region. A consensus in glycan processing was not detected at sites including N130 and N188 (**Figure 2E**), with factors such as differing variable loop length, surrounding glycan proximity, and local protein architecture modulating the processing state of glycans at these sites. Together, these examples illustrate the dynamic nature of variable loop glycans, wherein local structural constraints can reproducibly promote high-mannose glycosylation at certain positions (e.g., N156, N160), while at others, the processing state is less predictable and dependent on the surrounding glycan context. As glycans surrounding the V2 epitope can obstruct nAb binding, it is plausible that inter-strain heterogeneity in glycan processing could offer a route for glycan-mediated bnAb resistance.

### Conserved fine glycan processing defines the silent face of diverse HIV-1 strains

Across all glycan-dependent bnAb epitopes, the SF constitutes the most conserved epitope with respect to glycan processing (**Figure 2E-F**, bottom). This underscores the structural rigidity of this epitope across diverse strains and clades. Essential for native gp120 folding^76^, the N262 glycan site is one of the most highly conserved glycan sites on Env (>99%, **Figure 1C**), making it an attractive target for bnAbs. This site is most consistently occupied by M9 glycans (average 42.9%, **Figure 2E**) owing to its buried location at the gp120 protomer interface. The N262 site, along with N448, form primary carbohydrate contacts for SF bnAbs, including SF12^77^. Like N262, glycans occupying N448 are highly underprocessed. However, unlike N262, N448 glycosylation undergoes further mannose trimming, with M8 representing the most abundant glycan composition on average (33.1%). Though not explicitly an epitope for SF bnAbs, the N442 site, which is present in six of the panel Envs, is situated proximally to the primary bnAb epitope region. Interestingly, this site is highly occupied by oligomannose-type glycans, although the pattern of mannose trimming is reversed compared to N262 and N448, where M5 is most abundant (average 55.1%) and M9 is the least (average 0.5%). The presence of N442 and N448 is not mutually exclusive; the CH119, CE0217, and 25710 strains possess both sites. Presently, the extent to which N442 influences the ability of SF bnAbs to recognize their epitope is not fully understood.

The N295 site plays a prominent role in the recognition of both SF and V3 glycan bnAbs. In both instances, contacts between the CDRH3 loops and oligomannose-type glycans at N295 are established to support bnAb binding. A range of oligomannose glycoforms were detected at this site across all six of the N295-containing strains, with very low complex- and hybrid-type glycans present on average (0.8% and 1.8%, respectively). In CNE55, only 42.4% glycan occupancy was observed. Although SF bnAbs may not rely on N295 for high potency neutralization, this finding further emphasizes the principle that an encoded PNGS does not necessarily constitute efficient glycan attachment, and this observation could have wider implications for V3-targeted bnAb recognition.

### Strain-specific PNGS underoccupancy shapes divergent apex glycan signatures

The consensus glycosylation at N156 and N160 in **Figure 2F** shows that M5 is prevalent across most strains, with a median of 77.4% and 84.6%, respectively. As mentioned, previous studies have elucidated the preference of V2 apex bnAbs for small high mannose-type glycans at the trimer apex^73,74^. One of the twelve panel pseudoviruses, 398F1, deviated from this trend. Rather than the typical high-mannose phenotype that has been frequently observed in this study and others, a substantial proportion of complex-type glycans were detected at N160 (**Figure 4A**). The 398F1 strain is highly resistant to neutralization by V2 apex bnAbs (IC_50_ >25 µg/mL), distinguishing it from the other 11 strains in the panel. While this strain contains the E169 resistance signature^78^, we hypothesized that unfavorable glycans at N156 and/or N160 could be a contributing signature of resistance. Unlike any other Env strain in the panel, partial glycan occupancy at N156 (V1 loop) and N188 (V2 loop) was observed, averaging at 53.1% and 32.8%, respectively (**Figure 3**). However, V2 apex bnAbs typically mediate binding through contacts with N160 glycans as opposed to those at N156, so incomplete glycan occupancy at N156 likely does not directly impart neutralization resistance. Rather, this underoccupancy may allow for more advanced glycan processing, giving rise to the observed complex- (48.7%) and hybrid-type glycans (5.5%) at N160 through enhanced steric accessibility to glycan processing enzymes (**Figure 4B**), as has been observed before on soluble trimeric immunogens^31,43,79^. It is conceivable that such alterations in glycan processing at the apex could offer an additional molecular basis for 398F1 resistance to glycan-dependent V2 apex bnAbs, a feature not evident from the primary sequence alone. To further evidence the atypical nature of this glycosylation feature, N156, N160, and N188 on X2278, and N186 on CE0217 are fully occupied, giving rise to underprocessed high mannose-type glycans necessary for high affinity V2 apex bnAb binding. It is plausible that strain-specific differences in Env trimer dynamics could also contribute to the apex glycosylation observed on 398F1.

**Figure 4.**
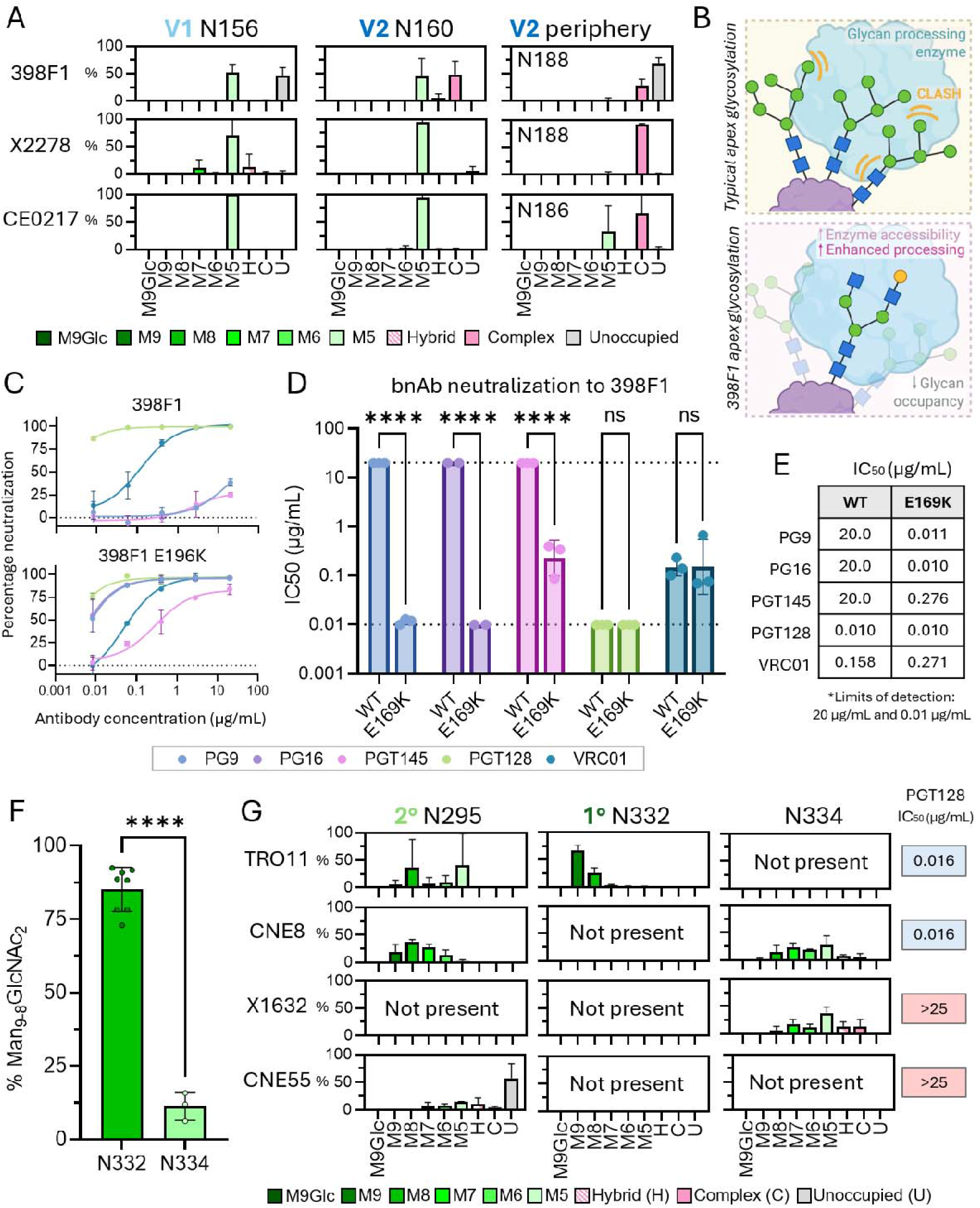
Strain-specific divergence in glycan epitope presentation at key mannose patch regions and the effect on *in vitro* virus neutralization. **A)** Difference in fine processing of key glycan sites implicated in recognition by V2 apex bnAbs. Percentage abundance of oligomannose, hybrid, complex-type glycans, or unoccupied glycan sites at N156, N160, and additional V2 glycan sites of the resistant 398F1 strain and sensitive CE0217 and X2278 strains. **B)** Schematic to demonstrate the principle of incomplete glycan occupancy and enhanced glycan processing. LC-MS data for 398F1 pseudovirus is used as motivation behind the schematic representation. **C)** Sensitivity of 398F1 to glycan-dependent V2 bnAbs (PG9, PG16, PGT145). The E169 resistance mutation is reverted to 169K to stratify any glycan-mediated contributions to V2 bnAb neutralization. V3-glycan (PGT128) and CD4bs bnAb (VRC01) used as controls. **D)** IC_50_ of different bnAbs to WT and E169K 398F1. The mean IC_50_ values are plotted for each bnAb tested (n=3, n=2 for PG16). The dotted lines refer to the limit of detection for the assay (20 μg/ml and 0.01 μg/ml). Two-way ANOVA was performed: ****, p < 0.0001. **E)** Table showing mean IC_50_ values presented in panel D. **F)** Percentage of Man_9-8_GlcNAc_2_ (M9-M8) glycosylation at N332 and N334. An unpaired T test was performed: ****, p < 0.0001. **G)** Difference in fine processing of key glycan sites implicated in recognition by V3 glycan bnAbs. Percentage abundance of oligomannose-type, hybrid-type, complex-type glycans, or unoccupied glycan sites at the secondary binding site glycan N295, and primary binding site glycan N332 or N334 for the highly sensitive TRO11 strain, selectively sensitive CNE8 strain, and broadly resistant X1632 and CNE55 strains. The geometric mean IC_50_ values of PGT128 against TRO11, CNE8, X1632 and CNE55 using data from the CATNAP database are indicated on the right side of the panel. Colors blue and red are used to indicate IC_50_ values of < 0.01 µg/mL and > 25 µg/mL, respectively. Error bars represent standard deviation in panels A, C, F and G. All statistical analyses performed using GraphPad Prism (version 10.3.1).

To investigate the extent to which these atypical N160 complex-type glycans contribute to bnAb resistance, we performed *in vitro* virus neutralization assays using wild-type (WT) 398F1 and an E169K reverted version to re-introduce the bnAb ‘sensitive’ sequence signature (**Figure 4C-E**). These analyses revealed that high potency (0.01 μg/mL) complete neutralization of 398F1 E169K could be achieved by PG9 and PG16, suggesting that these bnAbs are able to overcome glycan heterogeneity at N160. On the other hand, PGT145 could only achieve partial neutralization (between 79-94%) at the highest assayed concentration of 20 μg/ml (**Figure 4C**). So, it is plausible that a proportion of glycans at N160 on the 398F1 E169K mutant could be preventing a fraction of viruses from being neutralized *in vitro*. In all cases, a significant difference (p < 0.0001) in virus neutralization by V2-apex bnAbs could be measured between the WT and mutant pseudovirus as reflected by the relative IC_50_ values (**Figure 4D-E**). No significant differences in neutralization were detected for V3-glycan and CD4bs bnAbs PGT128 and VRC01, respectively (**Figure 4D**). Together, these results suggest that glycan heterogeneity at the trimer apex can differentially affect V2-apex bnAb neutralization and, in some contexts, may contribute to a fraction of viruses that are poorly neutralized or resistant.

### Elevated mannose trimming at N334 compromises the V3-glycan epitope

The network of glycans that comprise the high-mannose-rich IMP are sensitive to PNGS deletions. As one of the most highly targeted epitopes by bnAbs, and generally first to be targeted *in vivo*^80^, the integrity of the IMP is crucial for potent neutralization by V3-glycan bnAbs. The N332 glycan sits at the epicenter of the epitope targeted by V3-glycan bnAbs. Within the global 12-virus panel, eight strains contain the N332 glycan site, three contain the N334 glycan site, and one possesses neither. The N332 and N334 glycan sites are mutually exclusive^81,82^, and the presence of N334 is a common neutralization resistance feature. Across the global 12-virus panel, the V3-glycan bnAbs exhibit the largest range in IC_50_ values compared to any other bnAb epitope class, highlighting the fragility of this epitope^39^.

The consensus glycosylation for the N332 glycan site for the strains in the global 12-virus panel reveals a typical dominance of M9 and M8 glycans, with a median of 44.9% and 33.3%, respectively (**Figure 2F**). This, in addition to the presence of the intact ^324^GDIR^327^ motif, provides a molecular explanation for the high neutralization potency (IC_50_ <0.1 µg/mL) of 10-1074, PGT121, and PGT128 against TRO11, 25710, 398F1, X2278, BJOX002000, CE1176, CH119, and CE0217. On the other hand, glycans occupying the N334 glycan site display a significant increase in glycan processing compared to those at N332 (**Figure 4F**, p < 0.0001). Enhanced mannose trimming was observed at N334 on all three strains containing this site, giving rise to a range of M8-M5 glycans in addition to a small proportion of hybrid- and complex-type glycans. This glycan site shift reduces the abundance of oligomannose-type glycans with extended arms that are necessary for neutralization by several V3-glycan bnAb classes.

The TRO11 pseudovirus, which is potently neutralized by V3-glycan bnAbs, contains both N295 and N332. We detected glycans that are highly favorable for potent neutralization by glycan-dependent V3-glycan bnAbs (**Figure 4G**), with large underprocessed M9-M8 glycans present at N332 (average 66.6% and 26.3%, respectively) and a range of oligomannose-type glycans at N295, both of which are fully glycosylated.

The presence of N334 eliminates high potency neutralization by 2G12, 10-1074, and PGT121. However, PGT128 retains high neutralization potency against N334-containing strains including CNE8 and 246F3. Unlike members of the PGT121 class (PGT121-124, 10-1074), which exhibit a strong dependence on the N332 glycan for high affinity binding, PGT128 displays greater epitope flexibility; it can neutralize HIV-1 strains lacking the N332 site provided that the N295 glycan is present. To this end, we have identified a potential glycan-mediated mechanism of resistance to PGT128 that is not evident based on the sequence alone. The CNE8 and CNE55 Envs, members of subtype CRF01_AE, share 88% sequence identity. Both strains contain the N295 site but differ by the fact that CNE8 contains N334, yet CNE55 lacks this site. Unlike CNE8, the N295 site on CNE55 is predominantly unoccupied (average 57.5% unoccupancy, **Figure 4G**), which could provide a molecular basis for this strain’s resistance to neutralization by PGT128 (IC_50_ >25 µg/mL). In the case of X1632, the N332 and N295 sites are both absent, providing a clear and predictable explanation for the resistance phenotype. Similar to the observations with 398F1 and V2-apex bnAbs, these data highlight an additional layer of complexity in HIV-1 glycosylation. Glycan site conservation at the sequence level is important for bnAb recognition, but variation in glycan occupancy can also influence the antigenic properties of Env.

### Future perspectives

Over 250 published studies have utilized the global 12-virus panel to characterize monoclonal antibody responses^44^. Most of these antibodies directly contact or accommodate glycans to neutralize the virus, but due to the inherent flexibility of glycans are difficult to visualize using structural techniques. Unpicking the molecular determinants of bnAb neutralization resistance is increasingly necessary for successful HIV-1 vaccine development, particularly in the light of the results from the AMP trial^34^, where a significant population of circulating viruses exhibited pre-existing resistance to VRC01. Further, understanding how glycan heterogeneity can contribute to viral escape and shape viral rebound holds importance when considering the use of certain bnAbs in passive therapy and cure strategies. Our glycosylation analysis of the global 12-virus panel of pseudoviruses investigates glycan-mediated mechanisms of bnAb resistance and describes regions of the glycan shield that are robust, despite divergent protein sequences. This panel has permitted the standardized assessment of nAbs against diverse clades that represented the genetic diversity of circulating viruses at the time. This has allowed for the unification of neutralization data across different studies, to which these glycan analyses complement. Twelve years on since this virus panel was developed, and over twenty years since the HIV-1 isolates used in the panel were collected, an increasing trend in resistance by contemporary HIV-1 strains to clinically relevant bnAbs is being observed^35^. Understanding how Env is evolving over time and elucidating the mechanisms underpinning this will be useful for understanding the scale of Env diversity that needs to be overcome by a vaccine regimen. Determining the glycan-based resistance signatures emerging in contemporary strains forms an important component of this effort, particularly as glycan remodeling can alter bnAb epitopes without requiring extensive changes to the underlying protein sequence. Despite this increasing diversity, we identified several glycan sites that remain remarkably conserved and constitute key bnAb epitopes. Eliciting multiple bnAb lineages that neutralize several non-overlapping epitopes may be a key goal of vaccine design moving forward. So, focusing bnAb elicitation strategies on epitopes centered around conserved glycans, or by isolating or developing glycan-independent bnAbs, may offer a greater breadth of protection while presenting HIV-1 with a more constrained pathway for immune escape.

### Limitations of the study

The pseudoviruses analyzed in this study were produced in HEK293 cells. Whilst HEK293 cells faithfully represent the glycosylation of pseudoviruses routinely used in TZM-bl neutralization assays, some differences from viruses produced in primary PBMCs or *in vivo* are expected. Our previous analyses of BG505.T332N infectious molecular clones indicate that such differences are localized and have minimal impact on the overall glycan shield architecture^26^.

## STAR methods

## Experimental model and subject details

### Cell cultures

Female Human Embryonic Kidney (HEK) 293F cells were used to produce non-replicating HIV-1 pseudoviruses for glycan analysis by LC-MS. Cells were cultured in FreeStyle 293 Expression Medium (Thermo Fisher Scientific) and maintained at a density of 0.2-3 × 10^6^ cells/mL at 37°C, 8% CO_2_ and 125 rpm shaking. Cells were obtained from Thermo Fisher Scientific (catalogue number R79007).

Female HEK 293T cells were used to produce single-cycle HIV-1 pseudoviruses for *in vitro* virus neutralization assays. Cells were maintained in Dulbecco’s modified Eagle’s medium (Thermo Fisher Scientific) supplemented with 10% heat-inactivated fetal bovine serum (Thermo Fisher Scientific). Cultures were incubated at 37°C, 5% CO_2_ and cell monolayers were split 1:10 at confluence by treatment with 0.25% trypsin, 1 mM EDTA (Thermo Fisher Scientific). Cells were obtained from American Type Culture Collection (catalogue number CRL-3216).

### Env expression plasmids

Expression plasmids encoding HIV-1 Env were obtained from BEI Resources (National Institute of Allergy and Infectious Diseases (NIAID)), previously the NIH-AIDS Reagent Program, catalogue number #12670, “Global Panel of HIV-1 Env Reference Clones”. The HIV-1 Env molecular clones were constructed using the pcDNA3.1D/V5-His-TOPO^©^ expression vector and contain the ampicillin resistance gene.

The 398F1 E169K Env mutant was designed, synthesized, and cloned at GeneArt (Thermo Fisher Scientific). A GAA to AAG codon mutation was introduced into the original 398F1 Env gene, which corresponds to the E169K amino acid substitution, and the synthesized gene was cloned into the pcDNA3.1(+) expression plasmid.

## Method details

### Production of non-replicating HIV-1 pseudoviruses for glycan analysis

Prior to transfection, two solutions of 12.5 mL Opti-MEM (Thermo Fisher Scientific) medium were prepared. Packaging plasmids, pMDLg/pRRE and pRSV-Rev, and the Env expression plasmid were added to the first solution to give a final concentration of 310 μg/L at a ratio of 2:1:1, respectively. To the other solution, 1 mg/mL pH 7 polyethylenimine (PEI) max reagent was added to generate a ratio of 3:1 PEI max:plasmid DNA. Both solutions were combined and incubated for 30 minutes at room temperature. Cells were transfected at a density of 1 × 10^6^ cells/mL and incubated for 5 days at 37°C, 8% CO_2_ and 125 rpm shaking. Cells were centrifuged at 3041 × g for 30 minutes at 4°C and supernatant was applied to a 500 mL Stericup-HV sterile vacuum filtration system (Merck) with a pore size of 0.45 μm. Multiple independent transfections were performed for every pseudovirus strain, with each resulting sample preparation analyzed independently by mass spectrometry.

### Pseudovirus purification using *Galanthus nivalis* lectin (GNL)

Purification of non-replicating HIV-1 pseudoviruses for LC-MS analysis was performed by gravity-flow affinity chromatography using GNL-conjugated agarose (2BScientific). 5 mL GNL-conjugated agarose was added per 250 mL filtered supernatant and incubated overnight at 4°C. Supernatant was applied to a 50 mL glass gravity flow chromatography column (Bio-Rad) at a flow rate of 0.5-1 mL/min and washed with 150 mL PBS. Pseudoviruses were eluted from GNL using 50 mL 1 M methyl-α-D-mannopyranoside (Thermo Fisher Scientific) pH 3.5 (using acetic acid) and buffer exchanged into PBS using a Vivaspin column (molecular weight cut-off (MWCO) 100 kDa) (Sartorius) at 12°C. The GNL-conjugated agarose was washed with 150 mL PBS, followed by 150 mL 20% ethanol for storage.

### Site-specific compositional glycan analysis by LC-MS

An aliquot of 100 μg of purified HIV-1 pseudovirus (measured using UV-Vis spectroscopy where 1 Abs = 1 mg/mL) was denatured for 1h in 50 mM Tris/HCl, pH 8.0 containing 6 M of urea. Next, the sample was reduced and alkylated by adding 5 mM dithiothreitol (DTT) and 20 mM iodoacetamide (IAA) and incubated for 1h in the dark, followed by a 1h incubation with 20 mM DTT to eliminate residual IAA. The alkylated glycoproteins were buffer exchanged into 50 mM Tris/HCl, pH 8.0 using Vivaspin columns (10 kDa, Sartorius) and three aliquots were digested separately overnight using trypsin (Mass Spectrometry grade, Promega), chymotrypsin (Mass Spectrometry Grade, Promega) or alpha lytic protease (New England Biolabs) at a ratio of 1:30 (w/w) at 37°C. The next day, the peptides were dried and extracted using Oasis PRiME HLB 96-well μElution plate (Waters).

Prior to purification and desalting, the dried peptides were resuspended in 200 μL 0.1% trifluoroacetic acid (TFA). The Oasis PRiME HLB 96-well μElution plate was placed on a vacuum manifold, and the vacuum was set to 5” Hg. The HLB μElution wells were conditioned using 200 μL acetonitrile (ACN) then equilibrated with 200 μL 0.1% TFA. The vacuum was turned off, and the resuspended peptides were loaded into the conditioned wells. The vacuum was applied, starting at the lowest vacuum setting, and slowly increased to 5” Hg. The wells were washed with 800 μL 0.1% TFA, followed by 200 μL water to remove excess salts. The collection tray was placed in the vacuum manifold, and peptides were eluted using 80% ACN in 0.1% formic acid. The eluted peptides were dried prior to LC-MS analysis.

The dried peptides were re-suspended in 0.1% formic acid and analyzed by nanoLC-ESI MS with a Vanquish Neo (Thermo Fisher Scientific) system coupled to an Orbitrap Eclipse Tribrid mass spectrometer (Thermo Fisher Scientific) using stepped HCD fragmentation. Peptides were separated using a μPAC™ Neo HPLC Column (180 μm × 110 cm). A trapping column (PepMap 100 C18 3μM 75μM × 2cm) was used in line with the LC prior to separation with the analytical column. The LC conditions were as follows: 255-minute linear gradient consisting of 0–32% acetonitrile in 0.1% formic acid over 240 minutes, followed by 5 minutes of 76% acetonitrile in 0.1% formic acid. The flow rate was set to 300 nL/min and the spray voltage was set to 2.5 kV. The ion transfer tube temperature was set to 275 °C. The scan range was 350−2000 *m/z*. Stepped HCD collision energy was set to 15, 25 and 45% and the MS^2^ for each energy was combined. Precursor and fragment detection were performed using an Orbitrap at a resolution MS^1^= 120,000; MS^2^= 30,000. The AGC target for MS^1^ was set to standard and injection time set to auto which involves the system setting the two parameters to maximize sensitivity while maintaining cycle time.

### Site-specific assignment of glycan compositions

Glycopeptide fragmentation data were extracted from the raw file using Byos (Version 4.0; Protein Metrics Inc.). The glycopeptide fragmentation data were evaluated manually for each glycopeptide; the peptide was scored as true-positive when the correct b- and y-fragment ions were observed along with oxonium ions corresponding to the glycan identified. The MS data was searched using the Protein Metrics 305 N-glycan library with sulphated and phosphorylated glycans added manually. The relative amounts of each glycan at each site as well as the unoccupied proportion were determined by comparing the extracted chromatographic areas for different glycotypes with an identical peptide sequence. Identifications with a charge state of 1 were filtered out, all charge states for a single glycopeptide were summed. The precursor mass tolerance was set at 4 ppm and 10 ppm for fragments. A 1% false discovery rate (FDR) was applied. The relative amounts of each glycan at each site as well as the unoccupied proportion were determined by comparing the extracted ion chromatographic areas for different glycopeptides with an identical peptide sequence. Glycans were categorized according to the composition detected. HexNAc(2)Hex(9−3) was classified as M9 to M3. Any of these compositions that were detected with a fucose are classified as FM. HexNAc(3)Hex(5−6)Neu5Ac(0–4) was classified as Hybrid with HexNAc(3)Hex(5–6)Fuc(1)NeuAc(0–1) classified as Fhybrid. Complex-type glycans were classified according to the number of processed antenna and fucosylation. Complex glycans are categorized as HexNAc(3)(X), HexNAc(3)(F)(X), HexNAc(4)(X), HexNAc(4)(F)(X), HexNAc(5)(X), HexNAc(5)(F)(X), HexNAc(6+)(X) and HexNAc(6+)(F)(X). Core glycans are any glycan smaller than HexNAc(2)Hex(3).

### Site-specific glycan analysis of low abundance N-linked glycan sites using LC-MS

To obtain data for sites that frequently present low intensity glycopeptides, the glycans present on the glycopeptides were homogenized to boost the intensity of these peptides. This analysis can also be employed to obtain glycan analysis for sites that are situated in regions of high local glycan site density where the method described above cannot differentiate the glycans that are attached at >1 PNGS on a single peptide sequence. This analysis loses fine processing information but enables the ratio of high mannose: complex: unoccupied to be determined. The remaining glycopeptides from above were first digested with Endo H (New England Biolabs) to deplete oligomannose- and hybrid-type glycans and leave a single GlcNAc residue at the corresponding site. The reaction mixture was then dried completely and resuspended in a mixture containing 50 mM ammonium bicarbonate and Peptide:N-glycosidase F (PNGase F) (New England Biolabs) using only H_2_^18^O (Sigma-Aldrich) throughout. This second reaction cleaves the remaining complex-type glycans but leaves the GlcNAc residues remaining after Endo H cleavage intact. The use of H_2_^18^O in this reaction enables complex glycan sites to be differentiated from unoccupied glycan sites as the hydrolysis of the glycosidic bond by PNGaseF leaves a heavy oxygen isotope on the resulting aspartic acid residue. The resultant peptides were purified as outlined above and subjected to LC-MS. Instead of the extensive N-glycan library used above, two modifications were searched for: +203 Da corresponding to a single GlcNAc, and +3 Da corresponding to the ^18^O deamidation product of a complex glycan. Data acquisition and analysis was performed as above and the relative amounts of each glycoform were determined, including unoccupied peptides.

### *In vitro* virus neutralization assay

HIV-1 pseudotyped virus was generated in HEK-293T cells as previously described^83^. HEK-293T cells were transfected with plasmids expressing the HIV-1 virus backbone PSG3ΔEnv and full-length Env at a ratio of 2:1 using PEI (1mg/mL; 1:3 total DNA/PEI; Polysciences). Virus supernatants were harvested after 72h.

bnAb neutralizing activity was assessed using TZM-bl target cells that express CD4, CXCR4 and CCR5 receptors as described previously^16,84,85^. Antibodies were serially diluted and preincubated with virus for 1h before addition of TZM-bl cells (5,000 cells/well of a half-well plate). Each antibody concentration was tested in duplicates, and three independent assays were performed for each HIV-1 pseudotyped Env. Luminescence was quantified after 72h via lysis and addition of Bright-Glo luciferase substrate (Promega). ID_50_ was determined using nonlinear regression (GraphPad Prism version 10.3.1).

### Generating glycosylated Env models

To visualise the glycan shield of each Env strain, an AlphaFold 3 (Google DeepMind)^69^ model of the soluble trimer (inclusive of amino acids from position 40–664) was generated. Three copies of each chain were generated in the same job, and the resultant PDB was downloaded. A single chain was extracted as a PDB using PyMOL v2.5.2 and input into the Re-Glyco tool (https://glycoshape.org/reglyco) on the GlycoShape open access glycan structure database^70^. GlcNAc scanning was performed to identify all potential N-linked glycan sites, and a Man_5_GlcNAc_2_ glycan was modelled onto each identified site (GlyTouCan: G00028MO). The resultant fully glycosylated protomer was saved as a PDB and superimposed onto a single chain on the original AlphaFold 3-generated trimer in PyMOL. The fully glycosylated protomer was superimposed onto each chain of the trimer and extracted as an individual PDB so as to assemble three fully glycosylated protomers in a trimeric conformation. Glycans attached to each site were extracted and saved as individual PDBs. The unglycosylated, AlphaFold 3-generated soluble trimer and the individual glycans were visualized accordingly using ChimeraX v1.3.

### Quantification and statistical analysis

Mass spectrometry data was analyzed using Byos™ (Version 4.0), including the identification of glycopeptides and the XIC quantification of different glycoforms. All calculations and graphical representations of data were performed using GraphPad Prism version 10.3.1. Fluorescence intensities recorded for *in vitro* virus neutralization assays in Figure 4 were used to quantify the percent neutralization by each antibody dilution. Each antibody concentration was tested in duplicates, and the assay was independently repeated (n=3 for PG9, PGT145, PGT128, and VRC01; n=2 for PG16) to calculate the mean IC_50_ values for each antibody. In Figure 4D, a two-way ANOVA was performed where no significance is indicated by a p value >0.05 (denoted as not significant, “ns”), and **** indicates a p value of <0.0001. In Figure 4F, each plotted point represents the average percentage of M9-M8 glycosylation for each HIV-1 pseudotyped strain containing the indicated glycan site (N332, n=8; N334, n=3 where n represents the number of pseudotyped HIV-1 Env strains that contain the indicated PNGS). Bars represent the mean across the data points, and **** indicates a p value of <0.0001. In all figures, error bars represent the standard deviation (S.D.).

## Resource availability

### Lead contact

Requests for further information and resources should be directed to and will be fulfilled by the lead contact, Max Crispin.

### Materials availability

All unique reagents generated in this study are available from the lead contact with a completed materials transfer agreement.

### Data and code availability

Raw mass spectrometry files have been deposited on the MassIVE server (https://massive.ucsd.edu) under accession number MSV000102897 and are publicly available as of the date of publication. This paper does not report original code.

## Acknowledgements

We are very grateful to Michael Seaman and David Montefiori for critical reading of the manuscript. This work was supported by the Gates Foundation (INV-070116 to M.C.). A.M. was supported by the MRC-KCL Doctoral Training Partnership in Biomedical Sciences (MR/N013700/1). The following reagent was obtained through the NIH HIV Reagent Program, NIAID, NIH: Panel of Global Human Immunodeficiency Virus Type 1 (HIV-1) Env Clones, HRP-12670, contributed by Dr. David Montefiori.

## Author contributions

Conceptualization, M.L.N., J.D.A., and M.C.; Data Curation, M.L.N. and J.D.A.; Formal Analysis, M.L.N., J.D.A., and A.M.; Investigation, M.L.N., E.R.L., A.M., and N.E.J.; Resources, A.M. and K.J.D.; Supervision, J.D.A. and M.C.; Funding Acquisition, M.C., K.J.D.; Visualization, M.L.N.; Writing – original draft, M.L.N., J.D.A., and M.C. All authors contributed to reviewing and editing the manuscript.

## Supplemental information

Data S1. Glycan site alignment using Env sequences on the CATNAP database, related to Figure 1.

Data S2. Glycosylation of global 12-virus panel pseudovirions, related to Figures 2, 3, and 4.

## References

1. Cohen, K.W., De Rosa, S.C., Fulp, W.J., deCamp, A.C., Fiore-Gartland, A., Mahoney, C.R., Furth, S., Donahue, J., Whaley, R.E., Ballweber-Fleming, L., et al. (2023). A first-in-human germline-targeting HIV nanoparticle vaccine induced broad and publicly targeted helper T cell responses. Sci. Transl. Med. 15, eadf3309. 10.1126/scitranslmed.adf3309.

2. Steichen, J.M., Madden, P.J., Flynn, C.T., Phulera, S., Shil, M., Kalyuzhniy, O., Liguori, A., Kifude, C., Sewall, L.M., Cottrell, C.A., et al. (2026). Vaccination elicits HIV broadly neutralizing antibodies in primates. Nature 656, 723–733. 10.1038/s41586-026-10837-5.

3. Caniels, T.G., Prabhakaran, M., Ozorowski, G., MacPhee, K.J., Wu, W., van der Straten, K., Agrawal, S., Derking, R., Reiss, E.I.M.M., Millard, K., et al. (2025). Precise targeting of HIV broadly neutralizing antibody precursors in humans. Science 389, eadv5572. 10.1126/science.adv5572.

4. Steichen, J.M., Phung, I., Salcedo, E., Ozorowski, G., Willis, J.R., Baboo, S., Liguori, A., Cottrell, C.A., Torres, J.L., Madden, P.J., et al. (2024). Vaccine priming of rare HIV broadly neutralizing antibody precursors in nonhuman primates. Science 384, eadj8321. 10.1126/science.adj8321.

5. Moore, P.L., Williamson, C., and Morris, L. (2015). Virological features associated with the development of broadly neutralizing antibodies to HIV-1. Trends Microbiol. 23, 204–211. 10.1016/j.tim.2014.12.007.

6. Stephenson, K.E., Wagh, K., Korber, B., and Barouch, D.H. (2020). Vaccines and Broadly Neutralizing Antibodies for HIV-1 Prevention. Annu. Rev. Immunol. 38, 673– 703. 10.1146/annurev-immunol-080219-023629.

7. Hessell, A.J., Rakasz, E.G., Poignard, P., Hangartner, L., Landucci, G., Forthal, D.N., Koff, W.C., Watkins, D.I., and Burton, D.R. (2009). Broadly neutralizing human anti-HIV antibody 2G12 is effective in protection against mucosal SHIV challenge even at low serum neutralizing titers. PLoS Pathog. 5, e1000433. 10.1371/journal.ppat.1000433.

8. Saunders, K.O., Pegu, A., Georgiev, I.S., Zeng, M., Joyce, M.G., Yang, Z.-Y., Ko, S.-Y., Chen, X., Schmidt, S.D., Haase, A.T., et al. (2015). Sustained Delivery of a Broadly Neutralizing Antibody in Nonhuman Primates Confers Long-Term Protection against Simian/Human Immunodeficiency Virus Infection. J. Virol. 89, 5895–5903. 10.1128/JVI.00210-15.

9. Shingai, M., Donau, O.K., Plishka, R.J., Buckler-White, A., Mascola, J.R., Nabel, G.J., Nason, M.C., Montefiori, D., Moldt, B., Poignard, P., et al. (2014). Passive transfer of modest titers of potent and broadly neutralizing anti-HIV monoclonal antibodies block SHIV infection in macaques. J. Exp. Med. 211, 2061–2074. 10.1084/jem.20132494.

10. Andrabi, R., Bhiman, J.N., and Burton, D.R. (2018). Strategies for a multi-stage neutralizing antibody-based HIV vaccine. Curr. Opin. Immunol. 53, 143–151. 10.1016/j.coi.2018.04.025.

11. Steichen, J.M., Kulp, D.W., Tokatlian, T., Escolano, A., Dosenovic, P., Stanfield, R.L., McCoy, L.E., Ozorowski, G., Hu, X., Kalyuzhniy, O., et al. (2016). HIV Vaccine Design to Target Germline Precursors of Glycan-Dependent Broadly Neutralizing Antibodies. Immunity 45, 483–496. 10.1016/j.immuni.2016.08.016.

12. Caniels, T.G., Medina-Ramirez, M., Zhang, S., Kratochvil, S., Xian, Y., Koo, J.-H., Derking, R., Samsel, J., van Schooten, J., Pecetta, S., et al. (2024). Germline-targeting HIV vaccination induces neutralizing antibodies to the CD4 binding site. Sci. Immunol. 9, eadk9550. 10.1126/sciimmunol.adk9550.

13. Jardine, J.G., Kulp, D.W., Havenar-Daughton, C., Sarkar, A., Briney, B., Sok, D., Sesterhenn, F., Ereño-Orbea, J., Kalyuzhniy, O., Deresa, I., et al. (2016). HIV-1 broadly neutralizing antibody precursor B cells revealed by germline-targeting immunogen. Science 351, 1458–1463. 10.1126/science.aad9195.

14. Haynes, B.F., and Verkoczy, L. (2014). Host Controls of HIV Neutralizing Antibodies. Science 344, 588–589. 10.1126/science.1254990.

15. Verkoczy, L., and Diaz, M. (2014). Autoreactivity in HIV-1 broadly neutralizing antibodies. Curr. Opin. HIV AIDS 9, 224–234. 10.1097/COH.0000000000000049.

16. Walker, L.M., Huber, M., Doores, K.J., Falkowska, E., Pejchal, R., Julien, J.-P., Wang, S.-K., Ramos, A., Chan-Hui, P.-Y., Moyle, M., et al. (2011). Broad neutralization coverage of HIV by multiple highly potent antibodies. Nature 477, 466–470. 10.1038/nature10373.

17. West, A.P., Scharf, L., Scheid, J.F., Klein, F., Bjorkman, P.J., and Nussenzweig, M.C. (2014). Structural insights on the role of antibodies in HIV-1 vaccine and therapy. Cell 156, 633–648. 10.1016/j.cell.2014.01.052.

18. Burton, D.R., and Hangartner, L. (2016). Broadly Neutralizing Antibodies to HIV and Their Role in Vaccine Design. Annu. Rev. Immunol. 34, 635–659. 10.1146/annurev-immunol-041015-055515.

19. Doria-Rose, N.A., Schramm, C.A., Gorman, J., Moore, P.L., Bhiman, J.N., DeKosky, B.J., Ernandes, M.J., Georgiev, I.S., Kim, H.J., Pancera, M., et al. (2014). Developmental pathway for potent V1V2-directed HIV-neutralizing antibodies. Nature 509, 55–62. 10.1038/nature13036.

20. Briney, B.S., Willis, J.R., and Crowe, J.E. (2012). Human peripheral blood antibodies with long HCDR3s are established primarily at original recombination using a limited subset of germline genes. PLoS One 7, e36750. 10.1371/journal.pone.0036750.

21. Hwang, J.K., Wang, C., Du, Z., Meyers, R.M., Kepler, T.B., Neuberg, D., Kwong, P.D., Mascola, J.R., Joyce, M.G., Bonsignori, M., et al. (2017). Sequence intrinsic somatic mutation mechanisms contribute to affinity maturation of VRC01-class HIV-1 broadly neutralizing antibodies. Proc. Natl. Acad. Sci. U. S. A. 114, 8614–8619. 10.1073/pnas.1709203114.

22. Burton, D.R. (2019). Advancing an HIV vaccine; advancing vaccinology. Nat. Rev. Immunol. 19, 77–78. 10.1038/s41577-018-0103-6.

23. Sok, D., and Burton, D.R. (2018). Recent progress in broadly neutralizing antibodies to HIV. Nat. Immunol. 19, 1179–1188. 10.1038/s41590-018-0235-7.

24. Wibmer, C.K., Moore, P.L., and Morris, L. (2015). HIV broadly neutralizing antibody targets. Curr. Opin. HIV AIDS 10, 135–143. 10.1097/COH.0000000000000153.

25. Burton, D.R., Ahmed, R., Barouch, D.H., Butera, S.T., Crotty, S., Godzik, A., Kaufmann, D.E., McElrath, M.J., Nussenzweig, M.C., Pulendran, B., et al. (2012). A Blueprint for HIV Vaccine Discovery. Cell Host Microbe 12, 396–407. 10.1016/j.chom.2012.09.008.

26. Struwe, W.B., Chertova, E., Allen, J.D., Seabright, G.E., Watanabe, Y., Harvey, D.J., Medina-Ramirez, M., Roser, J.D., Smith, R., Westcott, D., et al. (2018). Site-Specific Glycosylation of Virion-Derived HIV-1 Env Is Mimicked by a Soluble Trimeric Immunogen. Cell Rep. 24, 1958–1966.e5. 10.1016/j.celrep.2018.07.080.

27. Cao, L., Pauthner, M., Andrabi, R., Rantalainen, K., Berndsen, Z., Diedrich, J.K., Menis, S., Sok, D., Bastidas, R., Park, S.-K.R., et al. (2018). Differential processing of HIV envelope glycans on the virus and soluble recombinant trimer. Nat. Commun. 9, 3693. 10.1038/s41467-018-06121-4.

28. Behrens, A.-J., and Crispin, M. (2017). Structural principles controlling HIV envelope glycosylation. Curr. Opin. Struct. Biol. 44, 125–133. 10.1016/j.sbi.2017.03.008.

29. Behrens, A.-J., Harvey, D.J., Milne, E., Cupo, A., Kumar, A., Zitzmann, N., Struwe, W.B., Moore, J.P., and Crispin, M. (2017). Molecular Architecture of the Cleavage-Dependent Mannose Patch on a Soluble HIV-1 Envelope Glycoprotein Trimer. J. Virol. 91, e01894–16. 10.1128/JVI.01894-16.

30. Pritchard, L.K., Spencer, D.I.R., Royle, L., Bonomelli, C., Seabright, G.E., Behrens, A.-J., Kulp, D.W., Menis, S., Krumm, S.A., Dunlop, D.C., et al. (2015). Glycan clustering stabilizes the mannose patch of HIV-1 and preserves vulnerability to broadly neutralizing antibodies. Nat. Commun. 6, 7479. 10.1038/ncomms8479.

31. Seabright, G.E., Cottrell, C.A., van Gils, M.J., D’addabbo, A., Harvey, D.J., Behrens, A.-J., Allen, J.D., Watanabe, Y., Scaringi, N., Polveroni, T.M., et al. (2020). Networks of HIV-1 Envelope Glycans Maintain Antibody Epitopes in the Face of Glycan Additions and Deletions. Structure 28, 897–909.e6. 10.1016/j.str.2020.04.022.

32. Seabright, G.E., Doores, K.J., Burton, D.R., and Crispin, M. (2019). Protein and Glycan Mimicry in HIV Vaccine Design. J. Mol. Biol. 431, 2223–2247. 10.1016/j.jmb.2019.04.016.

33. Crispin, M., and Doores, K.J. (2015). Targeting host-derived glycans on enveloped viruses for antibody-based vaccine design. Curr. Opin. Virol. 11, 63–69. 10.1016/j.coviro.2015.02.002.

34. Corey, L., Gilbert, P.B., Juraska, M., Montefiori, D.C., Morris, L., Karuna, S.T., Edupuganti, S., Mgodi, N.M., deCamp, A.C., Rudnicki, E., et al. (2021). Two Randomized Trials of Neutralizing Antibodies to Prevent HIV-1 Acquisition. New England Journal of Medicine 384, 1003–1014. 10.1056/NEJMoa2031738.

35. Mkhize, N.N., Yssel, A.E.J., Kaldine, H., van Dorsten, R.T., Woodward Davis, A.S., Beaume, N., Matten, D., Lambson, B., Modise, T., Kgagudi, P., et al. (2023). Neutralization profiles of HIV-1 viruses from the VRC01 Antibody Mediated Prevention (AMP) trials. PLoS Pathog. 19, e1011469. 10.1371/journal.ppat.1011469.

36. Wagh, K., Kreider, E.F., Li, Y., Barbian, H.J., Learn, G.H., Giorgi, E., Hraber, P.T., Decker, T.G., Smith, A.G., Gondim, M.V., et al. (2018). Completeness of HIV-1 Envelope Glycan Shield at Transmission Determines Neutralization Breadth. Cell Rep. 25, 893–908.e7. 10.1016/j.celrep.2018.09.087.

37. Bricault, C.A., Yusim, K., Seaman, M.S., Yoon, H., Theiler, J., Giorgi, E.E., Wagh, K., Theiler, M., Hraber, P., Macke, J.P., et al. (2019). HIV-1 Neutralizing Antibody Signatures and Application to Epitope-Targeted Vaccine Design. Cell Host Microbe 25, 59–72.e8. 10.1016/j.chom.2018.12.001.

38. Moore, P.L., Gray, E.S., Wibmer, C.K., Bhiman, J.N., Nonyane, M., Sheward, D.J., Hermanus, T., Bajimaya, S., Tumba, N.L., Abrahams, M.-R., et al. (2012). Evolution of an HIV glycan–dependent broadly neutralizing antibody epitope through immune escape. Nat. Med. 18, 1688–1692. 10.1038/nm.2985.

39. deCamp, A., Hraber, P., Bailer, R.T., Seaman, M.S., Ochsenbauer, C., Kappes, J., Gottardo, R., Edlefsen, P., Self, S., Tang, H., et al. (2014). Global panel of HIV-1 Env reference strains for standardized assessments of vaccine-elicited neutralizing antibodies. J. Virol. 88, 2489–2507. 10.1128/JVI.02853-13.

40. Panico, M., Bouché, L., Binet, D., O’Connor, M.-J., Rahman, D., Pang, P.-C., Canis, K., North, S.J., Desrosiers, R.C., Chertova, E., et al. (2016). Mapping the complete glycoproteome of virion-derived HIV-1 gp120 provides insights into broadly neutralizing antibody binding. Sci. Rep. 6, 32956. 10.1038/srep32956.

41. Go, E.P., Liao, H.X., Alam, S.M., Hua, D., Haynes, B.F., and Desaire, H. (2013). Characterization of host-cell line specific glycosylation profiles of early transmitted/founder HIV-1 gp120 envelope proteins. J. Proteome Res. 12, 1223– 1234. 10.1021/pr300870t.

42. Cao, L., Diedrich, J.K., Kulp, D.W., Pauthner, M., He, L., Park, S.-K.R., Sok, D., Su, C.Y., Delahunty, C.M., Menis, S., et al. (2017). Global site-specific N-glycosylation analysis of HIV envelope glycoprotein. Nat. Commun. 8, 14954. 10.1038/ncomms14954.

43. Behrens, A.J., Vasiljevic, S., Pritchard, L.K., Harvey, D.J., Andev, R.S., Krumm, S.A., Struwe, W.B., Cupo, A., Kumar, A., Zitzmann, N., et al. (2016). Composition and Antigenic Effects of Individual Glycan Sites of a Trimeric HIV-1 Envelope Glycoprotein. Cell Rep. 14, 2695–2706. 10.1016/j.celrep.2016.02.058.

44. Yoon, H., Macke, J., West, A.P., Foley, B., Bjorkman, P.J., Korber, B., and Yusim, K. (2015). CATNAP: a tool to compile, analyze and tally neutralizing antibody panels. Nucleic Acids Res. 43, W213–9. 10.1093/nar/gkv404.

45. Zhang, M., Gaschen, B., Blay, W., Foley, B., Haigwood, N., Kuiken, C., and Korber, B. (2004). Tracking global patterns of N-linked glycosylation site variation in highly variable viral glycoproteins: HIV, SIV, and HCV envelopes and influenza hemagglutinin. Glycobiology 14, 1229–1246. 10.1093/glycob/cwh106.

46. Marchitto, L., Wagh, K., Roark, R.S., Coleon, S., Li, H., Skelly, A.N., Hogarty, M.P., Habib, R., Ding, W., Ayyanathan, K., et al. (2026). Enhanced B cell priming induces broadly neutralizing HIV-1 apex antibodies. Nature, 656, 712–722. 10.1038/s41586-026-10838-4.

47. Coleon, S., Pathirage, R., Ding, W., Van Itallie, E., Mansouri, K., Marchitto, L., Cain, D.W., Lu, X., Kumari, V., Sutherland, L.L., et al. (2026). Rapid vaccine induction of macaque HIV-1 V2 Apex broadly neutralizing antibodies with immunogenetic signatures that are potentially translatable to humans. bioRxiv, 2026.05.27.728024. 10.64898/2026.05.27.728024.

48. Guenaga, J., Ádori, M., Bale, S., Phulera, S., Zygouras, I., Schleich, F.-A., Castro Dopico, X., Agrawal, S., Ota, M., Wilson, R., et al. (2026). Vaccination generates broadly cross-neutralizing antibodies to the HIV Env apex. Nature 654, 777–785. 10.1038/s41586-026-10429-3.

49. Habib, R., Roark, R.S., Li, H., Connell, A.J., Hogarty, M.P., Wagh, K., Wang, S., Marchitto, L., Skelly, A.N., Carey, J.W., et al. (2026). Env-antibody coevolution identifies B cell priming as the principal bottleneck to HIV V2 apex broadly neutralizing antibody development. Sci. Immunol. 11, eadz3933. 10.1126/sciimmunol.adz3933.

50. McCoy, L.E., van Gils, M.J., Ozorowski, G., Messmer, T., Briney, B., Voss, J.E., Kulp, D.W., Macauley, M.S., Sok, D., Pauthner, M., et al. (2016). Holes in the Glycan Shield of the Native HIV Envelope Are a Target of Trimer-Elicited Neutralizing Antibodies. Cell Rep. 16, 2327–2338. 10.1016/j.celrep.2016.07.074.

51. Derking, R., Allen, J.D., Cottrell, C.A., Sliepen, K., Seabright, G.E., Lee, W.-H., Aldon, Y., Rantalainen, K., Antanasijevic, A., Copps, J., et al. (2021). Enhancing glycan occupancy of soluble HIV-1 envelope trimers to mimic the native viral spike. Cell Rep. 35, 108933. 10.1016/j.celrep.2021.108933.

52. Freund, N.T., Wang, H., Scharf, L., Nogueira, L., Horwitz, J.A., Bar-On, Y., Golijanin, J., Sievers, S.A., Sok, D., Cai, H., et al. (2017). Coexistence of potent HIV-1 broadly neutralizing antibodies and antibody-sensitive viruses in a viremic controller. Sci. Transl. Med. 9, eaal2144. 10.1126/scitranslmed.aal2144.

53. Clark, M., Li, H., Martin, M., Evangelous, T.D., Gobeil, S.M.-C., Berry, M., Wagh, K., Giorgi, E.E., Hogarty, M.P., Zhao, C., et al. (2026). Humans and rhesus macaques share maturation pathways of HIV-1 envelope-reactive V3-glycan bnAb lineages. Sci. Transl. Med. 18, eaee9864. 10.1126/scitranslmed.aee9864.

54. Cottrell, C.A., Manne, K., Kong, R., Wang, S., Zhou, T., Chuang, G.-Y., Edwards, R.J., Henderson, R., Janowska, K., Kopp, M., et al. (2021). Structural basis of glycan276-dependent recognition by HIV-1 broadly neutralizing antibodies. Cell Rep. 37, 109922. 10.1016/j.celrep.2021.109922.

55. Briney, B., Sok, D., Jardine, J.G., Kulp, D.W., Skog, P., Menis, S., Jacak, R., Kalyuzhniy, O., de Val, N., Sesterhenn, F., et al. (2016). Tailored Immunogens Direct Affinity Maturation toward HIV Neutralizing Antibodies. Cell 166, 1459–1470.e11. 10.1016/j.cell.2016.08.005.

56. Tian, M., Cheng, C., Chen, X., Duan, H., Cheng, H.-L., Dao, M., Sheng, Z., Kimble, M., Wang, L., Lin, S., et al. (2016). Induction of HIV Neutralizing Antibody Lineages in Mice with Diverse Precursor Repertoires. Cell 166, 1471–1484.e18. 10.1016/j.cell.2016.07.029.

57. Dull, T., Zufferey, R., Kelly, M., Mandel, R.J., Nguyen, M., Trono, D., and Naldini, L. (1998). A third-generation lentivirus vector with a conditional packaging system. J. Virol. 72, 8463–8471. 10.1128/JVI.72.11.8463-8471.1998.

58. Stansell, E., Panico, M., Canis, K., Pang, P.-C., Bouché, L., Binet, D., O’Connor, M.-J., Chertova, E., Bess, J., Lifson, J.D., et al. (2015). Gp120 on HIV-1 Virions Lacks O-Linked Carbohydrate. PLoS One 10, e0124784. 10.1371/journal.pone.0124784.

59. Coss, K.P., Vasiljevic, S., Pritchard, L.K., Krumm, S.A., Glaze, M., Madzorera, S., Moore, P.L., Crispin, M., and Doores, K.J. (2016). HIV-1 Glycan Density Drives the Persistence of the Mannose Patch within an Infected Individual. J. Virol. 90, 11132– 11144. 10.1128/JVI.01542-16.

60. van Gils, M.J., Bunnik, E.M., Boeser-Nunnink, B.D., Burger, J.A., Terlouw-Klein, M., Verwer, N., and Schuitemaker, H. (2011). Longer V1V2 region with increased number of potential N-linked glycosylation sites in the HIV-1 envelope glycoprotein protects against HIV-specific neutralizing antibodies. J. Virol. 85, 6986–6995. 10.1128/JVI.00268-11.

61. Newby, M.L., Fogarty, C.A., Allen, J.D., Butler, J., Fadda, E., and Crispin, M. (2023). Variations within the Glycan Shield of SARS-CoV-2 Impact Viral Spike Dynamics. J. Mol. Biol. 435, 167928. 10.1016/j.jmb.2022.167928.

62. Pegg, C.L., Modhiran, N., Parry, R.H., Liang, B., Amarilla, A.A., Khromykh, A.A., Burr, L., Young, P.R., Chappell, K., Schulz, B.L., et al. (2024). The role of N-glycosylation in spike antigenicity for the SARS-CoV-2 gamma variant. Glycobiology 34, cwad097. 10.1093/glycob/cwad097.

63. Fadda, E., Singh, O., and Schulz, B.L. (2026). Heterogeneity of glycoproteins: Why does it matter and how to account for it. Curr. Opin. Struct. Biol. 99, 103296. 10.1016/J.SBI.2026.103296.

64. Williams, R.V., Huang, C., McDermott, C., Ahmed, T., Columbus, L., Moremen, K.W., Prestegard, J.H., and Amster, I.J. (2022). Site-to-site cross-talk in OST-B glycosylation of hCEACAM1-IgV. Proc. Natl. Acad. Sci. U. S. A. 119, e2202992119. 10.1073/pnas.2202992119.

65. Shrimal, S., and Gilmore, R. (2013). Glycosylation of closely spaced acceptor sites in human glycoproteins. J. Cell Sci. 126, 5513–5523. 10.1242/jcs.139584.

66. Xu, Y., Bailey, U.-M., Punyadeera, C., and Schulz, B.L. (2014). Identification of salivary N-glycoproteins and measurement of glycosylation site occupancy by boronate glycoprotein enrichment and liquid chromatography/electrospray ionization tandem mass spectrometry. Rapid Commun. Mass Spectrom. 28, 471–482. 10.1002/rcm.6806.

67. Crooks, E.T., Tong, T., Osawa, K., and Binley, J.M. (2011). Enzyme digests eliminate nonfunctional Env from HIV-1 particle surfaces, leaving native Env trimers intact and viral infectivity unaffected. J. Virol. 85, 5825–5839. 10.1128/JVI.00154-11.

68. Khaleque, M., Tropea, B., Singh, O., Elango, D., Liu, S., Allen, J.D., Newby, M.L., Sudol, A.S.L., Willcox, J., Butler, J., et al. (2025). Regulation of N-glycosylation efficiency by eukaryotic oligosaccharyltransferase. bioRxiv, 2025.09.06.674603. 10.1101/2025.09.06.674603.

69. Abramson, J., Adler, J., Dunger, J., Evans, R., Green, T., Pritzel, A., Ronneberger, O., Willmore, L., Ballard, A.J., Bambrick, J., et al. (2024). Accurate structure prediction of biomolecular interactions with AlphaFold 3. Nature 630, 493–500. 10.1038/s41586-024-07487-w.

70. Ives, C.M., Singh, O., D’Andrea, S., Fogarty, C.A., Harbison, A.M., Satheesan, A., Tropea, B., and Fadda, E. (2024). Restoring protein glycosylation with GlycoShape. Nat. Methods 21, 2117–2127. 10.1038/s41592-024-02464-7.

71. Caniels, T.G., Medina-Ramírez, M., Zhang, J., Sarkar, A., Kumar, S., LaBranche, A., Derking, R., Allen, J.D., Snitselaar, J.L., Capella-Pujol, J., et al. (2023). Germline-targeting HIV-1 Env vaccination induces VRC01-class antibodies with rare insertions. Cell Rep. Med. 4, 101003. 10.1016/j.xcrm.2023.101003.

72. Mason, R.D., Zhang, B., Morano, N.C., Shen, C.-H., McKee, K., Heimann, A., Du, R., Nazzari, A.F., Hodges, S., Kanai, T., et al. (2025). Structural development of the HIV-1 apex-directed PGT145-PGDM1400 antibody lineage. Cell Rep. 44, 115223. 10.1016/j.celrep.2024.115223.

73. Lee, J.H., Andrabi, R., Su, C.-Y., Yasmeen, A., Julien, J.-P., Kong, L., Wu, N.C., McBride, R., Sok, D., Pauthner, M., et al. (2017). A Broadly Neutralizing Antibody Targets the Dynamic HIV Envelope Trimer Apex via a Long, Rigidified, and Anionic β-Hairpin Structure. Immunity 46, 690–702. 10.1016/j.immuni.2017.03.017.

74. Doores, K.J., and Burton, D.R. (2010). Variable Loop Glycan Dependency of the Broad and Potent HIV-1-Neutralizing Antibodies PG9 and PG16. J. Virol. 84, 10510– 10521. 10.1128/JVI.00552-10.

75. Andrabi, R., Su, C.-Y., Liang, C.-H., Shivatare, S.S., Briney, B., Voss, J.E., Nawazi, S.K., Wu, C.-Y., Wong, C.-H., and Burton, D.R. (2017). Glycans Function as Anchors for Antibodies and Help Drive HIV Broadly Neutralizing Antibody Development. Immunity 47, 524–537.e3. 10.1016/j.immuni.2017.08.006.

76. Kong, L., Wilson, I.A., and Kwong, P.D. (2015). Crystal structure of a fully glycosylated HIV-1 gp120 core reveals a stabilizing role for the glycan at Asn262. Proteins 83, 590–596. 10.1002/prot.24747.

77. Schoofs, T., Barnes, C.O., Suh-Toma, N., Golijanin, J., Schommers, P., Gruell, H., West, A.P., Bach, F., Lee, Y.E., Nogueira, L., et al. (2019). Broad and Potent Neutralizing Antibodies Recognize the Silent Face of the HIV Envelope. Immunity 50, 1513–1529.e9. 10.1016/J.IMMUNI.2019.04.014.

78. Moore, P.L., Sheward, D., Nonyane, M., Ranchobe, N., Hermanus, T., Gray, E.S., Abdool Karim, S.S., Williamson, C., and Morris, L. (2013). Multiple pathways of escape from HIV broadly cross-neutralizing V2-dependent antibodies. J. Virol. 87, 4882–4894. 10.1128/JVI.03424-12.

79. Behrens, A.-J., Kumar, A., Medina-Ramirez, M., Cupo, A., Marshall, K., Cruz Portillo, V.M., Harvey, D.J., Ozorowski, G., Zitzmann, N., Wilson, I.A., et al. (2018). Integrity of Glycosylation Processing of a Glycan-Depleted Trimeric HIV-1 Immunogen Targeting Key B-Cell Lineages. J. Proteome Res. 17, 987–999. 10.1021/acs.jproteome.7b00639.

80. Bonsignori, M., Liao, H.-X., Gao, F., Williams, W.B., Alam, S.M., Montefiori, D.C., and Haynes, B.F. (2017). Antibody-virus co-evolution in HIV infection: paths for HIV vaccine development. Immunol. Rev. 275, 145–160. 10.1111/imr.12509.

81. Poon, A.F.Y., Lewis, F.I., Pond, S.L.K., and Frost, S.D.W. (2007). Evolutionary Interactions between N-Linked Glycosylation Sites in the HIV-1 Envelope. PLoS Comput. Biol. 3, e11. 10.1371/journal.pcbi.0030011.

82. Moyo, T., Kitchin, D., and Moore, P.L. (2020). Targeting the N332-supersite of the HIV-1 envelope for vaccine design. Expert Opin. Ther. Targets 24, 499–509. 10.1080/14728222.2020.1752183.

83. Montefiori, D.C. (2009). Measuring HIV Neutralization in a Luciferase Reporter Gene Assay. In Methods in Molecular Biology, 485, pp. 395–405. 10.1007/978-1-59745-170-3_26.

84. Krumm, S.A., Mohammed, H., Le, K.M., Crispin, M., Wrin, T., Poignard, P., Burton, D.R., and Doores, K.J. (2016). Mechanisms of escape from the PGT128 family of anti-HIV broadly neutralizing antibodies. Retrovirology 13, 8. 10.1186/s12977-016-0241-5.

85. Sarzotti-Kelsoe, M., Bailer, R.T., Turk, E., Lin, C., Bilska, M., Greene, K.M., Gao, H., Todd, C.A., Ozaki, D.A., Seaman, M.S., et al. (2014). Optimization and validation of the TZM-bl assay for standardized assessments of neutralizing antibodies against HIV-1. J. Immunol. Methods 409, 131–146. 10.1016/j.jim.2013.11.022.

